# Effort-based decision-making and gross motor performance: Are they linked?

**DOI:** 10.1101/2020.01.31.928366

**Authors:** Simone V. Gill, Samuel J. Abplanalp, Laura Keegan, Daniel Fulford

## Abstract

The purpose of this study was to investigate the relationship between effort-based decision making and gross motor performance. Effort-based decision making was measured using a modified version of the Effort Expenditure for Rewards Task in which participants pressed a button on a keyboard to fill a bar on a screen for a monetary reward. Gross motor performance was measured with a walking task in which participants matched their steps to the beat of an audio metronome. We hypothesized that increased effort during the effort-based decision making task would be paired with an increase in steps taken per minute during the gross motor task. However, the results of this study indicated no relationship between the effort-based decision making task and the gross motor task. Planning rather than decision-making may have been the cognitive construct that governed our gross motor task. These findings can be beneficial when thinking about potential interventions for populations who experience deficits in motor performance and cognition as well as for understanding the relationship between both cognitive and motor performance in healthy adults.

## Introduction

Countless activities necessitate intact cognitive and motor processes (1–3); for example, making it safely across a busy intersection requires deciding to cross during a gap in oncoming traffic and having the motor wherewithal to cross the street quickly. For impairments in cognition and motor performance, tasks requiring both processes pose a significant challenge to quality of life. On one hand, associations between cognitive and motor performance are plausible from both a neurological and conceptual standpoint. It is possible that aspects of cognition and motor performance are linked due to overlapping neurological substrates that are responsible for each area (4–7). For instance, the prefrontal cortex and the anterior cingulate gyrus have been implicated in decision-making and in monitoring actions (8). From a conceptual perspective, the stages involved in decision-making are intertwined with motor performance. Decision-making involves making a preference, executing an action, and experiencing an outcome (8), These steps suggest that cognitive (selection and planning) and motor (performing an action) processes (9) are inherent in decision-making, during which ongoing feedback guides action online (10–12).

On the other hand, the cognition involved in decision-making and motor performance may be separate (13), which is representative of the views taken via traditional models and theories (i.e., dualism) (14–17). The action execution step in decision-making may only refer to “action” as taking action or making a decision rather than executing a motor action. If so, then the neurological substrates implicated in decision-making may monitor taking action instead of gross motor execution (18,19).

Effort-based decision-making involves making a choice whether a given outcome (typically a reward) is worth the mental or physical effort required to obtain it (20). Previous studies have examined effort-based decision-making using the effort expenditure for rewards task (EEfRT) (21). The EEfRT is comprised of making a choice between either exerting physical effort for a monetary reward (i.e., money) via keyboard button pressing, or exerting no effort for less of a reward or no reward. The EEfRT has been used in studies of psychiatric populations, including people with schizophrenia and major depression, with results commonly showing that these groups demonstrate altered effort allocation by failing to make high-effort response choices that maximize reward. Prior research has hypothesized that deficits in effort-based decision-making may be due in part to motor impairments (22); however, no study has examined this association using reliable, validated, and sensitive assessments of motor performance.

The purpose of this study was to examine the relationship between effort-based decision-making and gross motor performance. We tested effort-based decision-making using a modified version of the EEfRT, and motor performance with a task that involved matching walking steps to the sound of an audio metronome (23). We hypothesized that a higher proportion of effortful trials chosen in the effort-based decision-making task would correspond with an increase in cadence (steps taken per minute) on the gross motor task.

## Method

Ethics Statement: The study and consent procedures were approved by the Boston University Institutional Review Board and conformed to the Declaration of Helsinki. Informed written and verbal consent was obtained from all participants before testing began.

### Participants

Fifty-nine participants responded to online or print flyers distributed in the Boston area or by word of mouth. Potential participants were screened and excluded if they endorsed any of the following: Below the age of 18 or above 65; had ever had a heart attack; ever been diagnosed with angina, asthma, cystic fibrosis, cardiovascular disease, bronchitis, obstructive lung disease, or a neurological or orthopedic condition; ever had a stroke; or if a physician had advised them against mild to moderate exercise. These exclusion criteria were in place because one of the primary tasks of the larger study included a vigorous exercise task (24).

### Metronome Task

Participants completed a gait task in which they were instructed to match their steps to an audio metronome beat as they walked along a 6.10m X .89m-long Protokinetics Gait Carpet (Protokinetics, LLC; Peekskill, NY, USA) Participants walked in 4 conditions: initial baseline, slow, fast, and final baseline. They first walked in an initial baseline condition that included 10 walking trials at their own pace (i.e., no metronome beat). For the metronome task, the average cadence (steps per minute) of each participant was calculated using measures from the initial baseline condition. Next, participants completed 10 trials in which the metronome beat was 15% slower than their baseline cadence and 10 in which the beat was 15% faster. Participants then completed a final baseline condition for 10 trials at their own pace without the metronome beat. In between these trials, the participants ran in 2 intermediately dispersed trials after each metronome condition at a self-selected pace (i.e., no metronome beat). Eight participants were excluded due to incomplete data from this task.

### Effort-based Decision-making Task

We used a modified version of the Effort-Expenditure for Rewards Task (21) as a measure of effort-based decision-making (25) First, participants filled up a bar on a computer screen by tapping a computer key as quickly as possible using their non-dominant pinky finger. This procedure was completed three times to obtain a baseline measure of button-pressing speed. Varying amounts of monetary reward requiring different levels of baseline effort are displayed across a series of trials. The reward magnitude was shown as a dollar amount (range: $1-$5.73; based on four bins: $1.25-$2.39, $2.40-$3.49, $3.50-$4.60, and >$4.60), and the required effort level is shown by the height of a vertical bar (20%, 50%, 80%, or 100% of the participant’s maximum baseline button-pressing rate). Presented alongside each effortful option is the option of receiving $1 for no effort. There were 44 trials. An example trial may be a choice presented to either exert 80% of baseline effort expenditure for $2.40, or no effort for $1. These choices were hypothetical given participants were not required to exert effort in the moment. After making all of their choices, in another set of trials participants were then asked to either perform their selected choices or to reverse their initial decision and choose the non-effortful option for $1. One participant was excluded due to accepting every effortful choice. We chose to exclude them as their responses would provide no variability when conducting multilevel models.

### Data Analysis

After excluding participants, we had a final sample size of (*N* = 50; see Table 1 for descriptive statistics). The first level of analysis examined bivariate correlations (Table 2) between variables of interest, including age, gender, condition (initial baseline, 15% slower, 15% faster), and proportion of effortful trials chosen from the effort-based decision-making task.

**Table 1.** Descriptive statistics of Demographic information, Cadence, and Effort-based decision making (*N* = 50)

|  | Mean (%) | Standard<br>Deviation | Range |
| --- | --- | --- | --- |
| Age | 23.74 | 8.45 | 18-65 |
| Gender |  |  |  |
| Female | 48.0 | - | - |
| Male | 52.0 | - | - |
| Race |  |  |  |
| Non-Hispanic Caucasian | 62.0 | - | - |
| Asian-American | 30.0 | - | - |
| African-American | 4.0 | - | - |
| Multiracial | 2.0 | - | - |
| Other not listed | 2.0 | - | - |
| Cadence* |  |  |  |
| Baseline | 107.20 | 7.18 | 92-132 |
| 15% slower | 91.12 | 6.43 | 76-112 |
| 15% faster | 122.56 | 8.19 | 104-152 |
| Effort based-decision making<sup>+</sup> | .71 | .12 | .43-.93 |

Notes: \*measured in steps per minute; <sup>+</sup>proportion of effortful trials chosen

**Table 2.** Bivariate correlations between demographic information, cadence, and effort-based decision-making (*N* = 50)

|  | 1. | 2. | 3. | 4. | 5. |
| --- | --- | --- | --- | --- | --- |
| 1. Age |  |  |  |  |  |
| 2. Gender | .24 |  |  |  |  |
| 3. Baseline cadence | .15 | -.33* |  |  |  |
| 4. 15% slower cadence | .14 | -.31* | .97** |  |  |
| 5. 15% faster cadence | .13 | -.37** | .98** | .97** |  |
| 6. Effort-based decision-making | .15 | -.19 | .27<sup>+</sup> | .25<sup>+</sup> | .24<sup>+</sup> |

Notes: \* $p < .05$ ; \*\* $p < .01$ ; <sup>+</sup> $p \leq .09$

Multilevel logistic models were used to examine associations between the outcome variable (i.e., proportion of effortful trial chosen) and within and between subject-level covariates. We selected covariates for inclusion in the multilevel models based on bivariate correlations and theoretical interest. Within-subject covariates included reward amount and effort level required stemming from the effort-based-decision-making task. Between-subject covariates included age, gender, initial baseline cadence, slow pace cadence (15% slower than the initial baseline cadence), fast pace cadence (15% faster than the initial baseline cadence).

## Results

### Bivariate Correlations

The within-subject variable (i.e., proportion of effortful trials chosen) was not associated with any between-subject variables, including sex and age; however, there were trend-level associations between proportion of effortful trials chosen and baseline cadence (*r* = .27, *p* = .06), 15% slower cadence (*r* = .24, *p* = .093), and 15% faster cadence (*r* = .25, *p* = .08; Table 2). Three within-subject variables were associated with one-another. Sex was negatively associated with all cadence variables, suggesting an association between identifying as female and a slower cadence in all walking conditions.

### Multilevel Models

In multilevel models, we included the binary outcome of effort or no effort selected across trials as the dependent variable. After running the baseline model, we then ran three models, each including reward amount and effort required as within-subject predictors (i.e., at Level 1), and age and sex as between-subject variables (i.e., at Level 2). Cadence levels (i.e., baseline, 15% slower pace, and 15% faster pace) served as additional between-subject variables in the three separate models. Each model also tested cross-level interactions, including the interaction between age and sex and effort level and reward amount. Cross-level interactions were also tested with all cadence variables, including the interactions between baseline, 15% slower pace, and 15% faster pace and effort level and reward amount.

As shown in Table 3, reward amount and effort required were significant within-subject predictors of effortful trials chosen in all models. No between-subject variables, including all cadence variables, were significant predictors of effortful trials chosen. Further, there were no significant cross-level interactions in predicting effortful trials chosen from the effort-based-decision-making task.

**Table 3.**
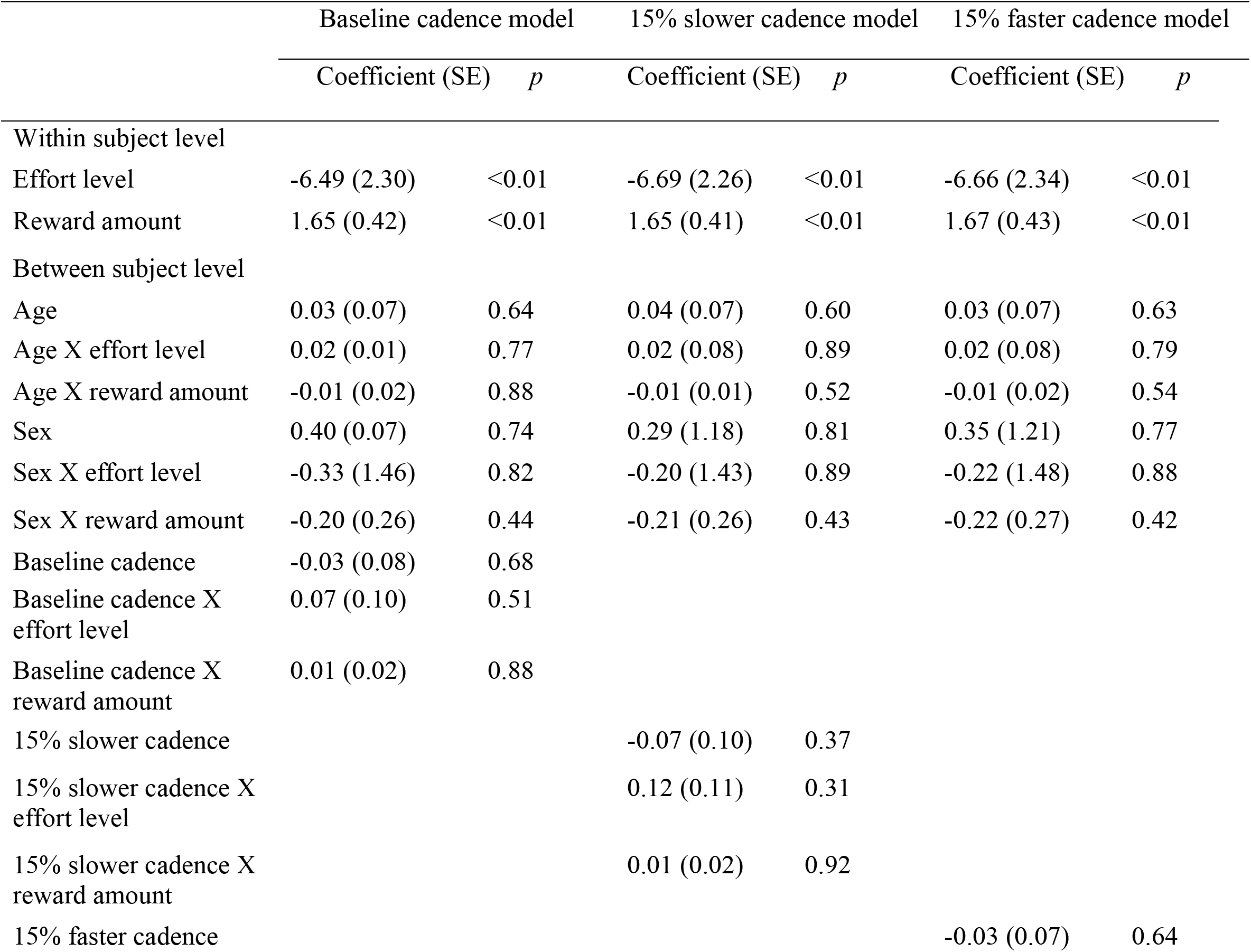
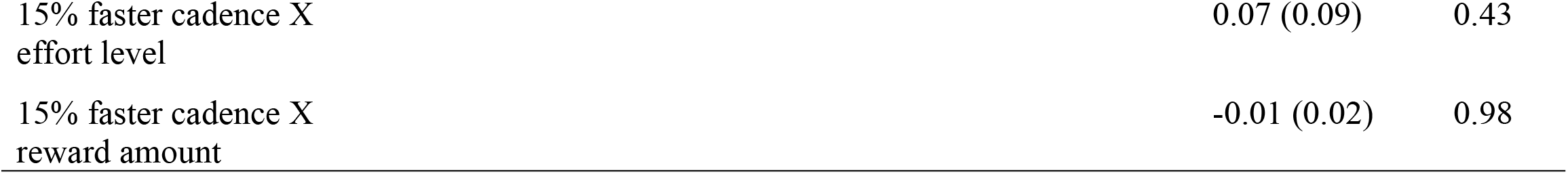
Within subject-level and between subject-level predictors of effortful choices (*N* = 50)

## Discussion

The purpose of this study was to investigate the relationship between effort-based decision-making and gross motor performance. Effort-based decision-making and gross motor performance were tested using an effort-based decision-making task and a walking task in which participants were asked to match their steps to an audio metronome. Inconsistent with our hypothesis, we identified no relationship between effort-based decision-making and cadence in any condition. Below, we propose three possibilities for our findings.

First, planning rather than decision-making may have been the cognitive construct that governed our gross motor task. Some researchers characterize planning as a part of decision-making in the context of gross motor performance (9,26,27). However, for gross motor performance, decision-making may align with deciding whether or not to perform a task (i.e., “taking action”) or selecting from multiple ways to perform a task (9) rather than monitoring the performance of the task (i.e., matching steps to a metronome beat) (23). Planning and decision-making may also have occurred serially rather than simultaneously (19,27,28) in our task: participants may have planned when to make initial heel contact prior to beginning the actual movement (14–17). Thus, the neurological substrates involved in deciding on versus monitoring gross motor performance may be different (18,29–31). Also, our findings would suggest that the action execution referred to in the steps of decision-making is separable from executing gross motor action (11,32).

Second, while the EEfRT (and similar tasks) is considered to reflect cognitive decision-making, some researchers have conceptualized it as a test of physical effort, given the involvement of fine motor skills in rapid button pressing (33). This task likely assesses the interaction of decision-making (as a cognitive process) and fine motor abilities. As such, the effort that a participant decides to exert may differ between fine and gross motor tasks; using a hand versus leg to complete a task may dictate the amount of effort exerted (34). For example, adults report differences in their perception of distances (35), jumping height (36), and objects (18) based on the effector (e.g., leg or hand). Future research needs to be done to disentangle the relationships among decision-making in the context of exerting effort to perform fine versus gross motor tasks.

Third, effort-based decision-making tasks offer rewards in relation to decisions that participants make, whereas our walking task provided no feedback about meeting the metronome pace. We intentionally omitted feedback to participants because we wanted to assess whether a relationship existed between effort-based decision-making and gross motor performance during a task that was purely focused on motor processes and because we thought that including the constraint to meet the metronome pace would transform walking into a complex, goal-oriented task (9,37) rather than the task of walking with no constraint (38,39). It could be that effort-based decision-making and motor performance are more likely to intersect when an error in achieving a goal is perceived as in the introduction of a perturbation (9) or in relation to the cost of making an error (27,40). Future studies will examine possible links between decision-making and gross motor performance in the presence of a reward and with the option to decide whether or not to complete the walking trial based on metronome pace.

Our findings are relevant to an understanding of cognitive and motor processes in healthy adults. These results are also important for helping psychiatric populations who experience challenges in cognition and motor performance. Our study challenges the assumption that interventions targeting decision-making would also lead to improved gross motor performance and vice-versa. Thus, separate intervention approaches may be needed to improve decision-making and gross motor performance to enhance quality of life. However, this assumption should be taken with caution because it is possible that this may not hold true in populations experiencing impairments in motor function, cognition, or both.

## Acknowledgments

We sincerely thank the participants and lab members for assistance with data collection and processing.

